# Comparative Analysis of Ecological and Germination Niches in Native and Invasive Populations of *Amaranthus albus*

**DOI:** 10.1101/2025.09.14.676160

**Authors:** Gal Rozenberg, Lior Blank, Yohay Carmel, Annie Euredjian, Mohsen B. Mesgaran

## Abstract

Invasive species serve as natural experiments to study adaptive evolution over contemporary time scales, as their native and invasive populations are exposed to different climatic settings and hence may experience strong selection pressures. In the life cycle of plants, germination stands as a cornerstone; thus, when invasive plant species encounter new surroundings, it is primarily expected that their germination niche will adapt to align with the thermal characteristics of the environment. However, adaptive responses in germination traits, specifically in cardinal temperatures, have not been explored. The objective of this study is to compare the *Amaranthus albus* (tumble pigweed) germination thermal niche and ecological niche between its native and invaded ranges. Considering geographic areas and climatic conditions, populations of *A. albus* were gathered from both cold and warm habitats in its native range (US) and its invaded range (Israel), with populations being at least 20 km apart from each other. First, populations were grown under similar conditions in a common garden experiment until seed production. The progeny seeds produced were then subjected to a set of germination trials under gradually changing temperature regimes. Cardinal temperatures and thermal niches for each population were estimated and analyzed in relation to the local climatic conditions of their respective habitats. Finally, a germination window was calculated to characterize the germination of a population throughout the year within its habitat. The invaded range studied here represents a subset of the species’ native range, with the *A. albus* establishing in similar climatic surroundings. Similarly, the germination niche for the native range was wider than for the invaded range. Thermal differentiation is evident in both native and invaded ranges. Populations from colder habitats exhibit lower base temperatures than those from warmer ones. This association was more evident for native populations. The remarkable variability observed within the germination pattern completely disappeared in the ‘germination window’ produced by those models, resulting in a similar pattern across the year for all populations. Our findings demonstrate the capability of species to adapt to new environmental conditions that may arise due to climatic changes. It emphasizes the role of cardinal temperatures, specifically the base temperature, as a potential adaptive characteristic.

## Introduction

Invasive species are more likely to establish in new areas when environmental conditions are similar to their native range (Thuiller et al. 2005). However, similarities among native and introduced ranges do not ensure a successful invasion. In their course of invasion, species must overcome a set of barriers acting on numerous scales, e.g., geographic, biotic, abiotic, dispersal, etc. (Richardson et al. 2000; Theoharides and Dukes 2007). The arrival of a species to a new region is often associated with a ‘founder effect’, where merely a small subset of the entire native population will be introduced, resulting in low genetic diversity in the invasive populations and thereby diminished trait variability (Dlugosch and Parker, 2008; Puillandre et al., 2008; Schrieber and Lachmuth, 2017). While low genetic variability could limit species spread, a considerable number of alien species have overcome this bottleneck via diverse biological, ecological, and evolutionary processes to become invasive, colonizing the introduced areas (Vilà et al. 2010; Schrieber and Lachmuth 2017). Novel environments are not necessarily hostile and might even be advantageous due to evolutionary changes, such as the evolution of increased competitive ability (Blossey and Notzold 1995), or as an ecological response, such as the enemy release hypothesis (Keane and Crawley 2002). Still, species need to alter their behavioral, physiological, or morphological traits to leverage these advantages. (Dlugosch and Parker, 2008).

A common framework to study species distribution in general, and invasive species in particular, is Ecological Niche Modelling (ENM). In essence, the set of environmental conditions where the species occurs delineates its ecological niche breadth (Sexton et al. 2017). Using ENMs, potential climatic conditions can be identified and compared across areas. The fundamental concept of ENMs, that is, utilizing a species niche to identify a suitable habitat of climate matching similarity, corresponds well with the prime principle of niche similarity in the invasion process (Peterson 2003). Therefore, ENMs are frequently employed to ascertain whether invasive species establish in the same conditions as in their native range, often referred to as ‘niche conservatism’, or spread into novel environments, known as niche shifts. However, it remains unclear which phenomenon is predominant, as both niche conservatism (Petitpierre et al. 2012; Liu et al. 2020) and niche shift (Atwater et al. 2018; Atwater and Barney 2021) find support in the literature. Empirical findings of the mechanisms that facilitate invasiveness were suggested to support the results obtained by niche modelling, i.e., niche shift or niche conservatism (Bates and Bertelsmeier 2021).

In their review, Donohue et al. (2010) described the profound impact of germination on a plant’s life. The transformation from seed to a plant marks a significant milestone, signifying a point of no return, governed by various factors like water availability and temperature. In irrigated land, where water is not considered a limited factor, temperature response constitutes a vital component. The conditions under which a seed germinates ultimately determine the environmental setting that the plant will encounter, suggesting that germination niches should correspond to individual ecological niches. Subsequently, germination traits are highly sensitive to selective forces, leading to rapid adaptations that may facilitate invasion (Donohue et al. 2005, 2010; Erfmeier and Bruelheide 2005; Hierro et al. 2009). Even small changes in temperature response can strongly influence whether a plant establishes, making it a useful way to detect early signs of adaptation. Despite the vast influence of germination on the plant life cycle, relatively few studies have evaluated the role of this trait in plant invasion or compared germination traits between native and invaded ranges (Leiblein-Wild et al. 2014; Gioria and Pyšek 2017).

Germination niche and cardinal temperatures, the range of temperatures within which germination occurs, may vary across populations originating in contrasting climatic conditions (Finch et al. 2019; Bürger et al. 2020; Marschner et al. 2024). Examining populations from distinct habitats in both native and invaded ranges can enhance our understanding of trait and niche shifts that may occur during invasion (Leiblein-Wild et al. 2014). Intriguingly, superior germination, which implies characteristics such as higher germination percentages or wider germination niche, is often reported for populations in their invaded rather than their native range, though other responses are reported (Gioria and Pyšek 2017). For example, the average germination niche of several *Ambrosia artemisiifolia* populations originating from its invaded range was wider compared to those originating from its native range (Leiblein-Wild et al. 2014). In contrast, *Verbascum thapsus* native populations exhibited a wider niche compared to invasive populations (Alba et al. 2016). Furthermore, the responses of germination traits to local ambient conditions within each range may exhibit diverging patterns (Hierro et al. 2009; Alba et al. 2016).

Originating from the southern parts of North America, *Amaranthus albus*, also known as *tumble pigweed*, has become a widespread invasive plant (Costea and Tardif, 2003). First documented in the region now known as Israel in 1879, its residency time is approximately 145 years. During this time period, *A. albus* has spread across agricultural fields and ruderal areas, spanning across a wide range of climatic conditions from arid to mesic Mediterranean climates (Dufour-Dror and Fragman-Sapir 2017; Gafni et al. 2023). *A. albus* is an annual plant that reproduces solely through seeds (Costea and Tardif, 2003); consequently, its germination traits are likely subject to selection (Alba et al. 2016). The germination cueing temperature is estimated to be 19°C for an invasive population (Cristaudo et al. 2014), and approximately 15°C (Steinmaus et al. 2000) for a native population, indicating potential variability in cardinal temperatures across different ranges or climatic conditions. The objective of this study is to compare the *A. albus* ecological niche and germination niche between its native and invaded ranges. Within this framework, we aim to 1) model and compare the climatic conditions occupied by *A. albus* in its native (US) and invaded (Israel) ranges, 2) evaluate our model results using empirical data from germination niche and, 3) test for evidence of thermal differentiation in germination characteristics between distinct climatic conditions of the native and invasive populations. Temperature responses of germination are quantified as an important trait in the context of climatic conditions, exploring how species may respond to these conditions.

The study hypothesizes the following: (1) The germination niche within each range is likely to reflect and indicate the climatic conditions where *A. albus* has established itself, as suggested by niche modeling. (2) As germination can influence the lifetime fitness of an individual, a parallel thermal differentiation in cardinal germination temperatures will be found in populations originating from distinct climates in both native and invaded ranges. Critical germination temperatures will follow similar patterns in both cold and warm habitats, regardless of whether the populations are native or invasive. However, (3) weaker patterns of climatic adaptation are expected in the invasive populations compared to native ones, because invasive populations have colonized their geographic ranges over shorter evolutionary timescales and are constrained by genetic variation.

## Materials and Methods

The ecological and germination niches of *A. albus* were compared between native and invaded ranges. While *A. albus* is a worldwide invasive plant, the invaded range considered in this study for both ecological niche and germination niche was restricted to the confined area of Israel. The ecological niche space was obtained via ecological niche modeling. Cardinal temperatures were obtained via the collection of populations from ‘warm’ and ‘cold’ habitats within each range (US and Israel).

### Ecological niche modeling

#### Occurrence data

Occurrence data for *A. albus* was downloaded from GBIF (the Global Biodiversity Information Facility). In order to improve the reliability and precision of the analysis, we addressed issues such as general coordinate validity, duplicate species coordinates, and the presence of country and province centroids. We used the *‘CoordinateCleaner’* package (Zizka et al. 2023). Common spatial and temporal inaccuracies were removed. In addition, multiple occurrences within a single cell were thinned down to just one for every pixel. Finally, 259 and 222 occurrences were retained for the native (US) and invaded (Israel) range, respectively.

#### Environmental data

Nineteen bioclimatic variables were obtained from the CHELSA climate dataset (version 2.1, available at http://chelsa-climate.org/) at 30 arc sec resolution (Karger et al. 2017). Bioclim rasters were resampled to a 5 km resolution to corresponding to the spatial resolution of occurrence in a large survey conducted a few years ago in the invaded range (Kadmon and Danin 1999) (Sillero and Barbosa 2021). Initially, variables were chosen to represent the mean and extreme climatic conditions (Mesgaran et al. 2014). However, additional screening was conducted in which variables with a high degree of multicollinearity (r > 0.7) were excluded (Sillero et al. 2021). Ultimately, four variables were considered to delineate the species’ ecological niche: precipitation of the driest and wettest quarter, as well as the mean temperature of the coldest and warmest quarter.

We defined the native range of *A. albus* by combining the species’ known native distribution as described by Govaerts et al. (2021), encompassing central and southern regions of the United States, as well as northern Mexico, with the World Wildlife Fund terrestrial ecoregions (Olson et al. 2001; Konowalik et al. 2017). This means we include all areas within terrestrial ecoregions where the species is found, regardless of whether they are originally mapped as a part of its native range. This approach disregards political boundaries, focusing instead on ecological relevance.

#### Ecological niche space

The obtained georeferenced occurrences and environmental data were utilized to construct the ecological niche space of *A. albus* in its native and invaded ranges. To compare the ecological niche of *A. albus* between its native and invaded range, we used the ‘ecospat’ package (Broennimann et al. 2023). The process involves the extraction of climatic conditions that prevail for each occurrence, followed by a principal component analysis (PCA) applied to both ranges simultaneously. By transforming correlated variables to uncorrelated principal components, PCA simplifies the dataset and addresses dimensionality and multicollinearity. The PCA scores were extracted to represent the position of each occurrence within the reduced environmental space. Subsequently, the ecological niche space was delineated and visualized by using these scores. Species occurrences were overlaid to visualize their distribution relative to environmental gradients. Finally, the ecological niche space was interpreted to compare species-environment relationships between the native and invaded ranges. The ecological niche dynamics were further validated using the ‘COUE’ framework (Guisan et al. 2014), which calculates four indices to compare invasive and native niches: (1) Centroid shift indicates changes in species’ preferences in a new environment compared to its native habitat; (2) Overlap assesses the proportion of shared environments in the native and invasive niches; (3) Unfilling measures how much niche space is unused by native species; and (4) Expansion quantifies the novel conditions that the species occupies within the invaded range.

### Germination, cardinal temperatures, and niche estimation

#### Seed collection

*A. albus* seeds collection was conducted in both the invaded (Israel) and native (US) ranges in accordance with the seed production season during the summer of 2021. A total of eight collection sites, four in the native and four in the invaded ranges, were selected to represent cold and warm climatic conditions within each respective range (Fig 1 & Table 1). Habitat distinction (warm vs. cold) was based on temperature rather than precipitation at the collection sites, since populations were sampled in irrigated land, minimizing differences in water availability. Species occurrence data extracted from GBIF, along with input from experts and extension officers, informed site selection. At each collection site, measures were taken to ensure that the sample represented the entire population adequately. Specifically, 30 mature plants were chosen haphazardly and harvested. Plants were selected adjacent to agricultural fields to mitigate potential confounding effects from agricultural practices such as irrigation and fertilization. The collected plants were bagged, dried, and then threshed to extract the seeds. Seeds from all plants at a particular location (hereafter, populations) were then pooled (i.e., a bulk seed collection).

**Table 1.** Climatic conditions obtained for the CHELSA climate dataset for the eight populations examined.

| Population | Range | Habitat | Average Annual Temperature (°C) | Temperature Range (°C) | Precipitation (mm) |
| --- | --- | --- | --- | --- | --- |
| Na1 | Native | Cold | 10.95 | 34.4 | 1020.7 |
| Na2 | Native | Cold | 10.15 | 38.4 | 388.1 |
| Na3 | Native | Warm | 16.95 | 29.8 | 554.9 |
| Na4 | Native | Warm | 18.05 | 31.4 | 202.7 |
| In1 | Invaded | Cold | 14.85 | 28.8 | 880.8 |
| In2 | Invaded | Cold | 19.95 | 19.2 | 782.1 |
| In3 | Invaded | Warm | 21.85 | 27.5 | 388.4 |
| In4 | invaded | Warm | 22.15 | 28.0 | 432.1 |

**Fig 1.**
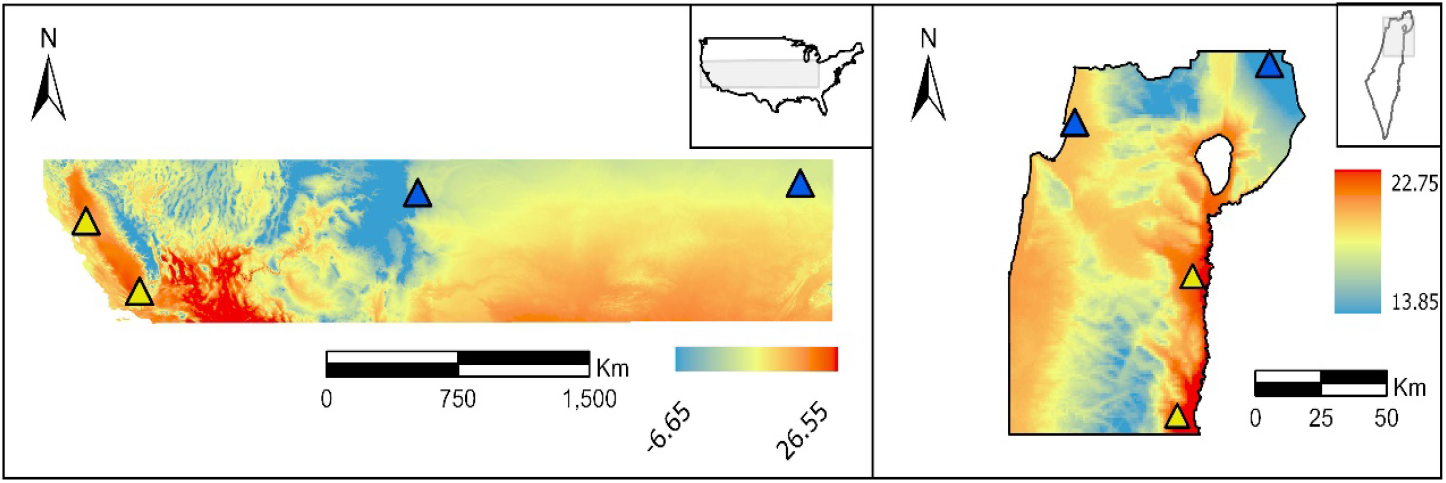
Map of native (A) and invaded (C) with a gray square designating the areas focused to illustrate mean annual temperature within each range (B and D). Populations considered as cold and warm habitats are represented by blue and yellow triangles, respectively.

#### Common garden experiments

To alleviate the potential impact of maternal effects (Gafni et al. 2024), two common garden experiments were conducted; one at UC Davis, USA and the other at Newe Ya’ar station, Israel. Seeds obtained for each population were sown in 30 pots, each filled with a commercial potting medium containing the slow-release fertilizer Osmocote®. Daily drip irrigation was applied to the pots. Upon reaching the 4-6 true leaf stage, seedlings were thinned, preserving one established seedling per pot. At the onset of flowering, pots originating from the same population were confined within a polyethylene cage as a preventive measure against cross-pollination. Following seed production, plants were harvested, dried at 40°C, and finally threshed. Produced seeds were stored in containers at room temperature for three months to reduce seed dormancy (Cristaudo et al. 2007). The simultaneous germination of all assays could not be completely coordinated due to technical issues across the two facilities. Thus, as a precautionary step, to minimize the potential effects arising from variations in storage duration (Gafni et al. 2024), seeds were subsequently kept at −20°C throughout the experiment, aiming to retain the seeds’ physiological condition unchanged during storage (Baskin and Baskin 2014).

#### Germination experiments

The germination experiment protocol was identical for all populations and temperatures. Before initiating assays at a designated temperature, seeds were moved from −20°C to 4°C for 24 hours to facilitate a gradual thawing process. Germination experiments took place across nine constant temperatures. Altogether, there were nine temperature treatments in this experiment, ranging from 5°C to 45°C with 5°C intervals. In the invaded range, a temperature treatment of 50°C was added. A wide spectrum of temperatures was essential, ensuring the estimation of both the base and ceiling temperatures. Fifty seeds were positioned within 9-cm-diameter Petri dishes on moistened Whatman® Grade 2 filter paper. Five replicates (i.e., Petri dishes), were used for each population × temperature combination. Dishes were sealed using parafilm® (Amcor, USA) and placed in a growth chamber (Conviron® Ltd., USA) set at a specific temperature under complete darkness. Dishes were counted daily under regular lighting for newly germinated seeds and moistened when required. Based on previous studies, the germination rate was expected to be high within the temperature range of 25-40°C. Therefore, to capture germination dynamics, dishes were monitored two to three times a day during the initial three to four days. After each count, dishes were placed in a random position in the growth chamber. The duration of assays varied, extending from 30 to 60 days contingent upon the timing of germination reaching saturation, with no seed germinations observed in the four days prior. To determine the viability of non-germinated seeds, dishes were incubated at 30°C for two weeks, followed by a ‘crush test’ (Borza *et al*., 2007; Baskin and Baskin 2014).

#### Seed germination modeling

Seed germination analysis was conducted in R (R Development Core Team, Vienna, Austria) using ‘drc’ (Ritz and Strebig 2016), ‘drcte’ (Onofri et al. 2023) and ‘aomisc’ (Onofri 2020a) packages. We calculated germination rate and fitted non-linear models to determine cardinal temperatures and thermal niche breadth for each population. First, the relation between cumulative germination and time was fitted via the nonparametric maximum likelihood estimator for time-to-event data for each dish in individual temperature employing *drmte()* function of *drcte* package (Onofri et al. 2023). The observed germination curves varied across our populations and temperature tests. To maintain methodological uniformity when fitting different models, the nonparametric approach was preferred due to its capacity to describe diverse shapes of time-to-event curves (Onofri et al. 2022; Marschner et al. 2024). Next, we calculated germination rate (GR) i.e., the reciprocal of the time until 20% of all seeds germinated (Keshtkar *et al*., 2021), using the *quantile* function of *drcte package* for the dish in each population × temperature combination. The 20^th^ percentile threshold was chosen because some populations failed to obtain higher cumulative germination in most temperatures. Subsequently, we regressed GR against temperature (T) for the estimation of three cardinal temperatures: base temperature, *T*_b_ indicating the lowest temperature at or below which germination does not occur; the optimal temperature *T*_O_, where germination rate peaks; and the ceiling temperature at or beyond which germination ceases. We applied the nonlinear beta regression (Yin *et al*., 1995) from the ‘aomisc’ package (Onofri 2020b) to describe the relation between *GR* and *T:*

Where *GR*_20_ is the germination rate for the 20^th^ percentile (fraction). *T* represents temperature, while *T*_*b*_, *T*_*o*_ and *T*_*c*_ denote the base, optimal, and ceiling temperatures, respectively. R_*max*_ indicates the maximum germination rate, attained at the optimal temperature (*T* = *T*_*o*_). The index *i* distinguishes individual populations analyzed, each characterized by its unique set of model parameters. Cardinal temperatures (model parameters) across populations within each species were fitted to data from all populations simultaneously for each range. The germination niche for the native and invaded ranges was calculated as the maximum ceiling temperature and the minimum base temperatures across all the populations in their respective range.

To compare thermal differentiation within the native and invaded range, the four populations collected from each range were categorized as originating from either ‘cold’ or ‘warm’ habitats (Table 1). Mean values from each habitat were obtained from their respective populations, and the models were compared.

#### Germination window

Germination threshold models were employed to predict the germination rate of populations at each month within their specific collection site, thereby generating each population ‘germination window’ (Filipe et al. 2023). Mean daily maximum temperature, averaged over the years 1981-2010, was obtained from the CHELSA climate dataset (version 2.1) at a resolution of 30 arc seconds (Karger et al. 2017). Monthly temperature data was then obtained for the different collection sites. Subsequently, the respective germination models for each collection site were applied to these monthly temperature datasets. This methodical approach facilitated an examination of germination patterns throughout the year for each population under consideration.

## Results

### Ecological niche space comparison

Principal Component Analysis effectively captured 78.0% of the variability inherent in the four bioclim variables used. Specifically, the first principal component (PC1) and second principal component (PC2) contributed 44.2% and 33.8% of the variance, respectively. PC1 primarily represents temperature, exhibiting negative high correlations with both temperature-related variables, whereas PC2 is mainly associated with precipitation variables (Fig. 1).

The ecological niche of the species within its native range encompasses a wider range of climatic conditions than in its invaded range. Despite the extensive fundamental niche observed within the native range, the species’ realized niche, delineating the specific climatic conditions it occupies, is considerably narrower, as further evidenced by the unfilling index of 40%. The realized niche of the *A. albus* is entirely confined within its native niche. This is further supported by niche dynamics indices, demonstrating perfect stability (=100%) and no expansion into new climates. The centroid shift indicates a slight transition towards warmer, drier conditions in the invaded range (Fig 2).

**Fig 2.**
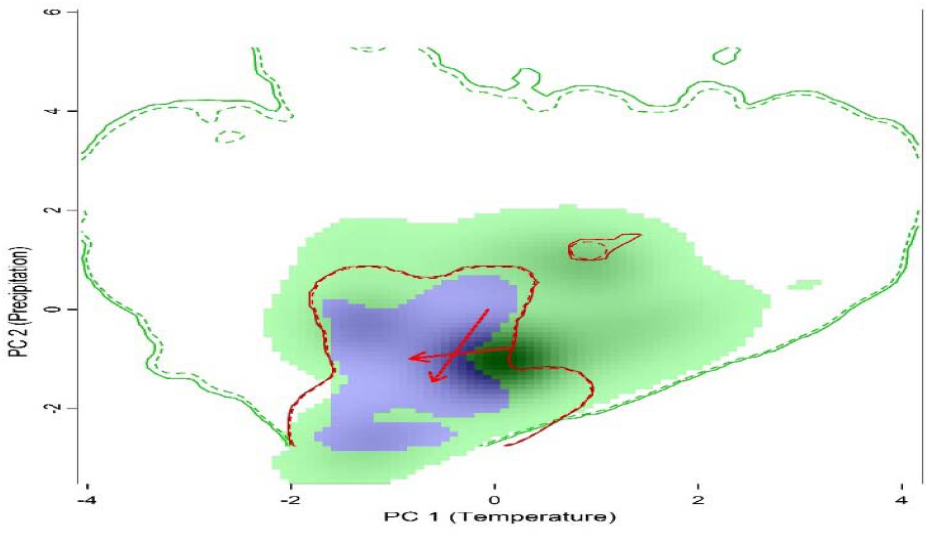
The ecological niche space of native (green) and invaded (red) ranges. Solid contour lines delineate the climatic conditions, in the native and invaded ranges, respectively. Niche stability is indicated in blue. Arrows illustrate the centroid shift from native to invasive distribution (continuous line) and between native and invaded extent (dashed line).

### Germination niche and cardinal temperature

The non-linear beta regression model accurately represents the dataset resulting in reliable estimates for cardinal temperatures (Fig. 3). This is evidenced by the small standard errors associated with these parameters, as detailed in Table 2. Maximum germination rate was generally higher in the invaded range, though the highest was obtained for Na3 in the native range. The lowest maximum GR was obtained for the two populations originating from cold habitats: Na1 and Na2.

**Table 2.** Estimated cardinal temperatures (*T_b_*: base temperature, *T_o_*: optimal temperature, *T_c_*: ceiling temperature) for different populations of *A. albus* within the native and invasive range. These estimates were obtained by fitting a beta function to germination rate calculated for time to 20% germination.

| Population | Range | Habitat | $T_b$ | $T_o$ | $T_c$ | Max GR |
| --- | --- | --- | --- | --- | --- | --- |
| Na1 | native | Cold | 14.4±1.53 | 35.7±1.57 | 40±0.03 | 0.6±0.09 |
| Na2 | native | Cold | 17.6±3.43 | 34.7±1.95 | 40±0.04 | 0.3±0.05 |
| Na3 | native | Warm | 19.9±2.35 | 37.6±0.38 | 40.1±0.03 | 1.5±0.25 |
| Na4 | native | Warm | 26.4±0.21 | 33.9±0.26 | 44.7±0.57 | 0.8±0.12 |
| In1 | invaded | Cold | 19.9±0.13 | 32.1±0.44 | 46.8±1.53 | 1.3±0.08 |
| In2 | invaded | Cold | 18.7±1.5 | 32.5±0.65 | 47.1±2.09 | 1.3±0.08 |
| In3 | invaded | Warm | 22.5±2.17 | 33.2±0.82 | 40.8±0.04 | 1.4±0.06 |
| In4 | invaded | Warm | 22.9±1.71 | 34.5±0.54 | 43.2±0.94 | 1.1±0.05 |

**Fig 3.**
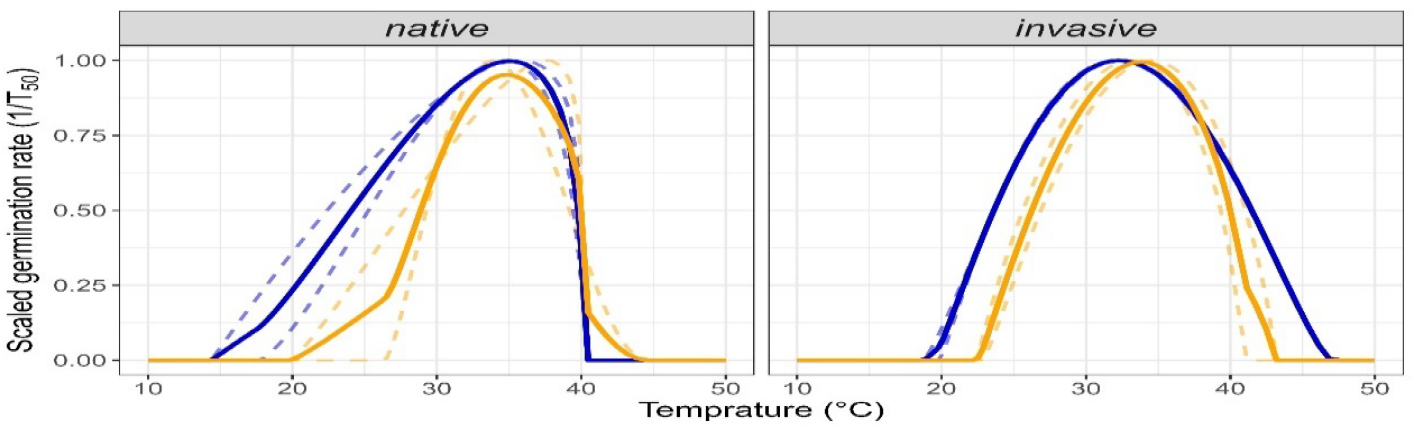
Fitted beta regression to germination rates for native and invasive populations originated from cold (blue) and warm (yellow) habitats. Solid line represents the mean of the populations in dashed lines.

The native and invaded ranges exhibited a broad spectrum of cardinal temperatures. In the native range, base germination exhibited the highest variability, ranging between the outmost population in the cold and warm habitats. Optimal and ceiling temperatures did not share the same variability. In the invaded range, base temperature showed lower variability but still exhibited a detectable three-degree difference between the Na2 and Na4 populations. Ceiling temperature also exhibited a wide range, spanning approximately six degrees.

Populations originating from cold habitats exhibited higher ceiling temperature. Germination niche for each range, that is, the difference between the maximum ceiling temperatures and the minimum base temperature, varied. The native germination niche spanned from 14.4±1.53°C to 44.7±0.57°C and was wider by about 2°C compared to the invaded range niche that spanned from 18.7±1.5°C to 47.1±2.09°C.

### Thermal differentiation

In the native range, the mean difference between the base temperature of ‘cold’ and ‘warm’ habitats is 7.1±3.14°C whereas in the invaded range, the difference is approximately 3.4± 2.23°C. Smaller negligible differences was found for the optimal temperatures of 0.5±1.8°C and 1.6±0.89°C the native and invaded range, respectively. Ceiling temperatures exhibited a 2.4±0.4°C difference for the native range while the invasive populations exhibited an opposing trend wherein cold populations were comprised of higher temperatures with a 5±1.95°C.

### Germination window

The annual germination window was similar across all populations (Fig. 4). In most cases, germination starts approximately in March, peaks in July and diminishes in October. Native populations exhibited a nearly indistinguishable pattern. However, the germination window for the invasive population was prolonged compared to the native populations. When populations were pooled according to habitat, (cold and warm) and range (native and invaded), no significant differences were found in the duration of the window.

**Fig 4.**
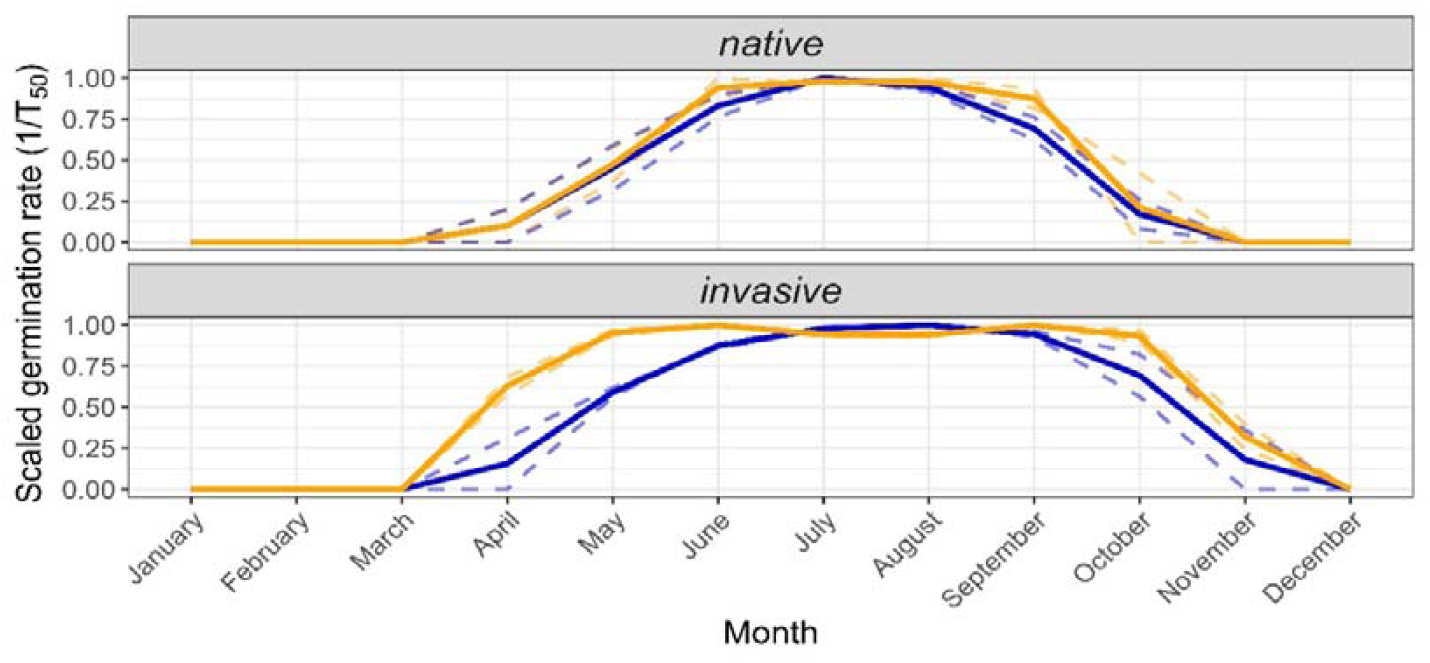
The germination window for native and invasive populations originated from cold (blue) and warm (yellow) habitats. Solid line represents the mean of the populations in dashed lines.

## Discussion

Given the crucial role of germination in the plant life cycle, the thermal response of invasive plants to their new environment would be expressed first and foremost in a modification of their germination niche. Despite its crucial role, germination traits are rarely compared between native and invaded ranges (Gioria and Pyšek 2017). Even fewer studies have utilized populations from various climatic conditions within each range (Hierro et al. 2009; Leiblein-Wild et al. 2014; Alba et al. 2016). In this study, we evaluated populations of *A. albus* from cold and warm habitats in their native and invaded range. We compared the ecological niche and germination profiles between native and invasive populations. The invaded range studied here, although representing a sharp climatic gradient from desert to Mediterranean climates (Rozenberg et al. under review), represents a subset of the species’ native range, with the *A. albus* conserving its ecological niche by establishing in similar climatic surroundings. Similarly, the germination niche for the native range was wider than for the invaded range. Thermal differentiation was observed in both the native and invaded ranges, wherein populations originating from cold habitats within their range had a lower base temperature than the respective populations from warm habitats, with the native population exhibiting a stronger association to local climatic conditions.

### Ecological and germination niche

*A. albus* established in similar climatic conditions within the native and the studied invaded range, and thus conserved its niche. Overall, most invasive plants were found to conserve their ecological niche (Petitpierre et al. 2012; Liu et al. 2020) though there are contradictory studies supporting niche shift (Atwater et al. 2018; Atwater and Barney 2021); in some cases, small-scale differences in local climate or soil conditions could allow plants to occupy slightly different niches that are not apparent at the larger scale of analysis. The result of this study is unsurprising, as the climatic range in the invasive area studied here is completely contained within the range of climatic conditions in the native range of *A. albus* (Bates and Bertelsmeier 2021). The studied invaded range is indeed smaller by orders of magnitudes compared to the native range and characterized by arid to mesic Mediterranean climates, which also comprise the native range in addition to others (Fig. 1). The climatic conditions within the invaded range studied here matched those of the native, likely facilitating the dispersal of *A. albus* during its course of invasion (Thuiller et al. 2005).

Consistent with the observed pattern for the ecological niche, wherein the invasive niche space was encapsulated within the native niche space, a similar trend was detected for the germination niche width. The invasive germination niche width was found to be narrower and mostly confined within the broader span of the native germination range, supporting our first hypothesis. This is the first study that compares the ecological niche with the germination niche, specifically between native and invaded ranges. Generally, the germination niche was suggested to correspond to the species ecological niche, yet studies evaluating this relationship reported contradictory results. For example, the germination pattern of a species was correlated with their respective ecological niche by Brändle et al. (2003), whereas Thompson and Ceriani (2003) reported no such association for Herbaceous species in the UK. This discrepancy may be explained by the limited number of populations and variations in ecotypes utilized to estimate the species’ germination niche width (Donohue et al. 2010; Finch et al. 2019). Indeed, in this study we observed large intraspecific trait variation in cardinal temperatures among populations. Therefore, collecting populations from diverse climatic conditions within each specified range should strengthen our findings.

Post-introduction adaptation in invasive species was previously reported in light of niche conservatism and suggested to facilitate the invasion process (Alexander 2013) (Sherpa et al. 2019). Ecotype variation and its correlation to local climatic conditions may stem from diverse mechanisms such as phenotypic plasticity, i.e., the ability of an individual to respond to its surroundings, and transgenerational maternal effects (Alba et al. 2016; Matzrafi et al. 2020), which occur when the environment experienced by a mother influences the phenotype or characteristics of her offspring. This study accounted for both by obtaining the germination niche using progeny seeds grown in a common garden experiment under a shared environment. Local adaptation is proposed as the source of variation and thermal differentiation. Local adaptations are a common phenomenon in both native and invaded ranges (Oduor *et al*., 2016). They occur at large scales, as well as in small areas, typically across steep environmental gradients (Erfmeier et al. 2011). As *A. albus* reproduces only by seeds (Costea and Tardif, 2003), its germination characteristics are under strong selection pressure (Donohue 2002; Verdú and Traveset 2005; Alba et al. 2016).

Trait variation may demonstrate both similar and diverging patterns across climatic clines in their native and invaded ranges (van Boheemen et al. 2019). In agreement with our second hypothesis, we found a parallel thermal differentiation in cardinal temperatures, specifically for base temperatures, in populations originating from distinct climates in both ranges; optimal and ceiling temperatures did not share a similar pattern. While other studies reported re-establishment of climatic clines for other traits (Montague et al. 2008), including germination traits (Hernández et al. 2019), this is the first time it has been reported for base temperature, to the best of our knowledge. Adaptations in base temperature are suggested to be valuable, as they facilitate a competitive advantage in resource acquisition (Gardarin et al. 2010; Gioria et al. 2018). In this study, populations originating from cold habitats demonstrated lower base temperature than populations from warm habitats in both the native- and invaded ranges. Although this finding appears straightforward, and in line with other studies (Fletcher et al. 2020), other reports suggest that populations from cold habitats have higher base temperatures (Alba et al. 2016; Bürger et al. 2020). Presumably, higher base temperatures facilitate delayed emergence, serving to avoid potential frosting events. Germination response to temperate conditions may possibly be climate- and taxon-dependent (Rosbakh and Poschlod 2015).

A comprehensive meta-analysis revealed that invasive species exhibit local adaptation to the same extent and intensity as native species (Oduor *et al*., 2016). Yet few studies focused on adaptive response in germination characteristics. For example, germination traits were correlated with local climatic conditions for *Verbascum Thapsus* within its native but not its invaded range (Alba et al. 2016). In our study, the difference in base temperatures between populations originating from cold and warm habitats was more pronounced within their native range compared to the invasive range, indicating stronger local adaptation as suggested in our third hypothesis. The limited geographical span and climatic conditions in the invaded range compared to the native range are acknowledged as potential contributors to the observed ‘weaker’ local adaptation. However, examining the ‘extreme’ populations from the US and Israel reveals a consistent difference of about eight degrees in the mean annual temperature between cold and warm habitats within each geographical range. Yet, the base temperatures’ difference in the two climatic habitats was twice as high in the native range. Other factors, such as low genetic variation due to the ‘founder effect’ (Dlugosch and Parker 2008a) and residency time (Gruntman and Segev 2024), hinder the ability of populations to adapt locally in their invaded range. The 145 years of residency time within the invaded range under study should be a sufficient period for the species to adapt to local conditions (Whitney and Gabler 2008; Colautti and Barrett 2013; Clements and Jones 2021) and it is plausible that, as time progresses, differences in the invaded range could become increasingly distinct across different habitats.

Interestingly, the maximum germination rate differed between cold and warm habitats but only in the native range. Notably, low germination rates were observed for the two populations originating from cold habitats. In contrast, the populations originating from warm habitats exhibited high germination rates, comparable to all populations from the invaded range. Indeed, the invaded range studied here is a subset of the conditions within the native range mostly characterized by higher annual temperatures. Even the ‘cold’ habitat in the invaded range demonstrated a higher annual temperature compared to the warm habitats in the native range (Table 1). This may indicate the species general preferences to warm climates, as also suggested by the optimal temperature, which was above 30°C for all the populations in this study and other reports for this species (Steckel et al. 2004; Cristaudo et al. 2007; Gafni et al. 2024). It was suggested that invasive species exhibit higher germination rates in their invaded range (Gioria and Pyšek 2017). In this study, germination rates do not appear to be influenced by the populations’ range, i.e., native or invaded, but rather it is the population’s climatic conditions that matter. This highlights the importance of not only sampling in both native and invaded ranges, but also adequately representing the various climatic conditions within each range for a valid comparison.

The germination window, which is the duration in which a population is able to germinate according to its cardinal temperatures and *in-situ* temperate conditions, was similar in both the native and invaded ranges, lasting seven months starting in March. The marked variability observed within the germination pattern obtained via the models (Fig. 3) disappeared in the ‘germination window’ (Fig. 4) produced by those models. Apparently, all populations exhibited a generally similar temporal pattern of germination. This similarity was more pronounced in the native range, reinforcing our third hypothesis regarding weaker patterns of climatic adaptation in the invasive populations. The prolonged germination observed after the summer peak is less probable but still possible. *A. albus* germination depends on soil moisture, but in the invaded range, as well as the warm habitats in the native range, the soil dries out completely by the end of the summer. In addition, summer irrigated crops, which provide a favorable landscape for this species (Gafni et al. 2023), are harvested and irrigation ceases. The germination window can be improved by using a hydrothermal model that accounts for temperature and water availability, incorporating the required water potential with precipitation and irrigation schedules in the fields. Furthermore, high temperatures peaked at mid-summer are likely to facilitate seed dormancy (Batlla and Benech-Arnold 2015), as seed dormancy can act as a risk-reducing strategy to allow populations to avoid germinating in less favorable temperatures. The seed dormancy may last more than three months (Gafni et al. 2024) with highest dormancy relief acquired after six months (Cristaudo et al. 2007). This duration of seed dormancy generally corresponds with approximately five months where germination does not occur.

The invaded range studied here was limited, and thus did not fully capture the capacity of *A. albus* to undergo niche shifts. Further ecological niche modeling may include additional regions to which this species was introduced that are outside the geographical scope of this study. Niche conservatism was reported for another Amaranth species originating from North America, *Amaranthus retroflexus*, whose invaded range comprised of wide-ranging climatic conditions (Petitpierre et al. 2012). In this study, populations were collected to reflect the diverse climatic conditions within each range. The climatic conditions in the invaded range correspond mostly to the warm part of the native range. Thus, in some cases, the ‘cold habitats’ in the invaded range are sometimes warmer than the warm habitats of the native range. For example, one invasive population classified as a cold habitat had a higher annual temperature than the two populations classified as warm in their native range. Additionally, the relatively small sample size hinders the possibility of conducting a more robust statistical comparison across ranges and habitats. Future work can also genetically compare native and invasive populations to identify the ‘founding’ population(s) and the role of continuous gene flow in the adaptation observed in *A. albus*, similar to another invasive plant in Israel that originated from the US (Hübner et al. 2022). Despite these limitations, our findings provide important insights into the crucial role of germination adaptations in plants. Comparing species responses in native and invaded ranges can indicate contemporary evolution and the capacity of species to withstand climatic changes (Colautti and Lau 2015). The remarkable contrast between the observed variation in base temperatures and the similarity in germination window highlight the species capacity for adaptation to local climatic conditions. The fact that this trend was observed in both native and invaded ranges highlights the species ability to adapt to climatic changes.

## Acknowledgments

We gratefully acknowledge the support of the BSF (US-Israel Binational Science Foundation) grant 132-2101, awarded to Lior Blank and Mohsen B. Mesgaran. Special thanks to Roni Gafni from Newe Ya’ar Research Center for her invaluable support and insights. We also thank Talya Avraham for her meticulous work in the germination experiments. Additionally, we appreciate the collaboration of Itay Shulner and Evyatar Asaf from Ran Lati’s lab at Newe Ya’ar Research Center for their contribution to the field experiments. Our thanks extend to Hanan Eizenberg and Maor Matzrafi from Newe Ya’ar Research Center for their knowledge and facilities, which greatly enhanced our research.

We gratefully acknowledge the support of the BSF (US-Israel Binational Science Foundation) grant 132-2101, awarded to Lior Blank and Mohsen B. Mesgaran.

## Notes

### Competing Interest Statement

The authors have declared no competing interest.

